# Testosterone and cortisol are negatively associated with ritualized bonding behavior in male macaques

**DOI:** 10.1101/765123

**Authors:** Alan V. Rincon, Michael Heistermann, Oliver Schülke, Julia Ostner

**Affiliations:** Department of Behavioral Ecology, Johann-Friedrich-Blumenbach Institute for Zoology and Anthropology, University of Goettingen, Goettingen, Germany; Leibniz ScienceCampus Primate Cognition, Goettingen, Germany; Endocrinology Laboratory, German Primate Center, Leibniz Institute for Primate Research, Goettingen, Germany; Research Group Social Evolution in Primates, German Primate Center, Leibniz Institute for Primate Research, Goettingen, Germany

**Keywords:** Testosterone, cortisol, steroid/peptide theory, social bond, triadic male-infant-male interaction, Barbary macaque

## Abstract

Neuroendocrine research on the formation of social bonds has primarily focused on the role of nonapeptides. However, steroid hormones often act simultaneously to either inhibit or facilitate bonding. Testosterone is proposed to mediate a trade-off between male mating effort and nurturing behavior; therefore, low levels are predicted during periods of nurturing infant care and social bonding. In species where social bonding and support regulates hypothalamic-pituitary-adrenal axis activity, we also expect glucocorticoid levels to be low during bonding periods. We investigated how immunoreactive urinary testosterone (iuT) and cortisol (iuC) were related to triadic male-infant-male interactions – a ritualized male bonding behavior – as well as infant care in male Barbary macaques (*Macaca sylvanus*). We collected >3000 hours of behavioral observation data during full-day focal animal follows from 14 adult males and quantified iuT and iuC from 650 urine samples. As predicted, both iuT and iuC were negatively correlated with rates of triadic interactions within-subjects in the hours preceding urination. We found no relationship between iuT and iuC with triadic interactions between-subjects. Infant care was weakly positively correlated to iuT and iuC within-subjects, but not between-subjects. The observed negative relationship between iuT and triadic interactions may be beneficial to lower competitive tendencies between adult males and to not inhibit bond formation. Lowered iuC could reflect increased bonding and perceived social support as triadic interactions predict future coalition formation in this species. Additionally, lowered iuC may be reflective of buffered tensions between males. The positive relationship of iuT and iuC with infant care suggests that the handling of infants in may be less nurturing but rather protective or competitive in this species. Measuring steroid hormones in relation to bonding and nurturing can help us interpret behaviors within the ecological contexts that they occur.

## 1 Introduction

Steroid hormones and behavior are intricately linked and often affect each other in a reciprocal manner (Goymann, 2009). Stable differences in baseline hormone levels could result in stable differences in behavioral profiles across individuals. Additionally, hormone levels can change dynamically within individuals to help them respond adaptively to a changing social environment. Distinguishing whether hormone-behavior interactions operate at the within- or between-subjects level (or both) can help us understand behavioral variation among individuals and behavioral flexibility within individuals (Hau and Goymann, 2015).

The challenge hypothesis (Goymann et al., 2019; Wingfield et al., 1990), proposes that high testosterone levels (regulated by the hypothalamic-pituitary-gonadal axis; HPG) promotes male mating effort and inhibits paternal care. Across vertebrates, males exhibit elevated testosterone levels when competing for mates, such as during the mating season or territorial defense, and low testosterone levels during periods of paternal care (Grebe et al., 2019; Hirschenhauser and Oliveira, 2006; Lynn, 2008; Muller, 2017; Wingfield et al., 1990). Testosterone potentially inhibits paternal care as experimentally increasing testosterone levels reduces paternal behavior in various bird species (Lynn, 2008), including increases within an individual’s natural range (Goymann and Flores, 2017). However, testosterone levels are positively related to paternal care when care is competitive in nature and concerns offspring defense from conspecifics (Moore et al., 2019; Muller, 2017). The steroid peptide theory of social bonds (S/P theory) builds on the challenge hypothesis to resolve this inconsistency in how testosterone relates to paternal care by proposing that high testosterone levels inhibit nurturing behaviors and intimacy in general rather than paternal care per se (van Anders et al., 2011).

Beyond mating effort and nurturing care, testosterone is relevant to bonding between same-sex adults. Same-sex adult bonds are usually characterized by affiliative contact and mutual social support (Massen et al., 2010; Ostner and Schülke, 2018) and share many qualities with nurturing infant care. Consequently, the S/P theory proposes that low levels of testosterone are ideal for bond formation to not inhibit the bonding process (van Anders et al., 2011). Consistent with this prediction, low testosterone has been linked to meat-sharing in male chimpanzees (*Pan troglodytes*: Sobolewski et al., 2012), a behavior implicated in bond formation (Wittig et al., 2014). Similarly, low basal levels of testosterone, and decreased levels from baseline, were associated with feelings of closeness following a friendship formation task in same-sex adult human dyads (Ketay et al., 2017).

Glucocorticoid steroids (primarily corticosterone in birds and rodents and cortisol in other mammals) are also involved in regulating animal social behavior, including in contexts relevant to nurture and social bonds. Glucocorticoid release is regulated by the hypothalamic-pituitary-adrenal (HPA) axis and is responsive to actual or perceived stressors (Sapolsky, 2002). The presence of or interaction with bonded partners during a potentially stressful event often attenuates HPA axis activity, a phenomenon known as social buffering (Cohen and Wills, 1985; Hostinar et al., 2014). Being well socially integrated or being able to count on others for social support may help to regulate HPA axis activity even in the immediate absence of stressors (Rosal et al., 2004; Wittig et al., 2016). As with testosterone, cortisol levels also decline in humans after increasing feelings of personal closeness, and individuals with low baseline cortisol had *partners* that desired to be closer to them (Ketay et al., 2017). Thus, mutually lowered cortisol and testosterone may have synergistic facilitative effects in the context of same-sex bonding.

Similarly to testosterone, a male’s glucocorticoid response to infant care may be dependent on social context. Within nurturing contexts, infant care appears to be negatively related to glucocorticoid levels. For example, cortisol levels decline following father-offspring interactions under non-stressful conditions in humans (Gettler et al., 2011b; Storey et al., 2011). Similarly, male marmosets (*Callithrix kuhlii*) that frequently carried infants had significantly lower glucocorticoid levels than males that carried infants less frequently (Nunes et al., 2001). However, glucocorticoid levels may be positively related to paternal care under more stressful contexts. For example, cortisol levels increase in response to infant cries in men (Fleming et al., 2002). In chacma baboons (*Papio ursinus*), males whose infants were at immediate risk of infanticide showed significant increases in glucocorticoid levels compared to a stable period, and had overall higher levels than males whose infants were not at risk (Cheney et al., 2015).

The aim of this study was to investigate how urinary testosterone and cortisol relate to same-sex bonding and nurturing behaviors in male Barbary macaques (*Macaca sylvanus*). Male Barbary macaques form stable social bonds with other males (Young et al., 2014b) that physiologically buffer bonded males during stressful situations (Young et al., 2014a). Two types of interactions are relevant to our aims: triadic male-infant-male interactions (hereafter triadic interactions) and infant care. Triadic interactions are ritualistic and involve two adult males sitting in body contact with an infant or young juvenile in-between them, exchanging affiliative facial signals and often lifting up the immature or inspecting its genitals (Deag and Crook, 1971; Paul et al., 1996). These interactions are of short duration, occur primarily during the non-mating season, predict coalition formation in the following mating season, and have been implicated in bond maintenance (Berghänel et al., 2011; Kalbitz et al., 2017). Triadic interactions may also function in ‘agonistic buffering’, where the presence of the infant allows two males to approach each other in a non-threatening context (Deag and Crook, 1971; Paul et al., 1996). Given the potential bonding and/or agonistic buffering function of triadic interactions, we predicted a negative correlation with both urinary testosterone and cortisol levels.

In addition to performing triadic interactions, male Barbary macaques also care for infants and yearlings on a dyadic level (in the absence of other males) by huddling, carrying, and grooming with them, sometimes for prolonged periods of time (Whitten, 1987). The exact function of male infant care in this species remains ambiguous. Males generally prefer to care for infants based on their probability of paternity inferred from their previous mating success with the mother (Kubenova et al., 2019), although their preferred infant is not always their genetic offspring (Ménard et al., 2001). One possible explanation is that males care for infants in a nurturing context. Another possibility is that infant care may be considered part of mating effort to increase a male’s mating success with the mother on the following mating season (Ménard et al., 2001). A previous study on this species suggested that handling infants might be stressful as it was associated with elevated fecal glucocorticoid levels (Henkel et al., 2010). Depending on the function of infant care, we may expect different relationships between this behavior with testosterone and cortisol. If infant care is nurturing in nature, we predict a negative relationship with testosterone (van Anders et al., 2011; Wingfield et al., 1990) and cortisol (Nunes et al., 2001; Storey et al., 2011). If the function of infant care is protective, or if infants are used as tools for mating effort, we expect a positive relationship between this behavior and testosterone (Muller, 2017; van Anders et al., 2011) and cortisol (Cheney et al., 2015).

Finally, as steroids can influence behavior within- or between-subjects we explicitly differentiate between both types of effects in our analyses. We predicted that the directionality of results would be the same whether effects are within- or between-subjects and that both types of effects would be found.

## 2 Materials and methods

### 2.1 Study site and animals

Study subjects belonged to one group of Barbary macaques (group C) living in semi-free ranging conditions within 14.5 ha of enclosed forest at Affenberg Salem, Germany (de Turckheim and Merz, 1984). Monkeys were provisioned once a day with fruits, vegetables, grains and had *ad libitum* access to water and monkey chow. Data were collected from 31 March to 26 October 2016 during one non-mating season. The study group consisted of 14 adult males (7 to 25 years old), 20 adult females (5 to 27 years old), 2 large subadult males (6 years old) and 19 immatures (up to 4 years old, 10 females, 11 males, including 2 male and 1 female yearlings, and one newborn male). The oldest adult male died during the study period on 20 August 2016. All group members could be individually recognized by observers though a combination of unique characteristics including body size, coat color and condition, facial spots and scars, gait and identification tattoos on the inner thigh.

### 2.2 Behavioral data collection and dominance hierarchy

Behavioral data were collected from 14 adult males using continuous focal animal sampling (Martin and Bateson, 2007). Focal males were followed continuously for an entire observation day (8-13.5 hours per day; total = 3289 hours, mean ± standard deviation = 235 ± 19 hours per individual; Table 1). The occurrence of all social interactions, including the identity of social partner(s) were recorded. We defined infant care as the time males spent in body contact with infants including huddling, carrying, and grooming, with ‘infants’ defined as < 1.5 years old (i.e. newborns and yearlings, N = 4). As infant care can occur for prolonged periods of time, we recorded the duration of this behavior. We defined triadic interactions as those that occurred between two adult males and any immature individual (0-4 years old, N = 19; see introduction for a description). Although triadic interactions can develop from a dyadic male-infant situation in some cases, they also develop when a male without an infant is approached by a male carrying an infant or by two males spontaneously converging on the same infant without prior male-infant interactions. In addition, the group of immatures involved in either of the two immature-related behaviors differed, making the two behaviors rather independent. As triadic interactions are often short in duration, we recorded the frequency of this behavior.

**Table 1:** Descriptive information per study subject for the study period. Behavior rates shown here were calculated based on all observation days for descriptive purposes. Behavior rates used in analyses were calculated individually for each urine sample and hormone excretion window.

| Male ID | Observation days | Observation hours | Dominance rank (norm David's score) | Triadic interactions (per obs. hr.) | Infant care (% obs. time) | Age (years) | Testosterone samples (N) | Cortisol samples (N) |
| --- | --- | --- | --- | --- | --- | --- | --- | --- |
| W3 | 19 | 238 | 14.6 | 0.81 | 3.32 | 12 | 13 | 24 |
| U2 | 20 | 243 | 13.1 | 0.45 | 1.16 | 13 | 42 | 42 |
| M3 | 21 | 235 | 12.6 | 0.06 | 2.16 | 21 | 19 | 20 |
| T3 | 21 | 242 | 11.1 | 0.51 | 2.42 | 14 | 42 | 56 |
| M4 | 21 | 247 | 10.9 | 0.04 | 0.07 | 21 | 43 | 49 |
| W7 | 19 | 224 | 10.3 | 0.31 | 0.64 | 12 | 40 | 44 |
| B5 | 20 | 238 | 7.9 | 1.80 | 6.65 | 7 | 55 | 66 |
| B6 | 21 | 247 | 7.3 | 0.65 | 2.14 | 7 | 57 | 61 |
| B7 | 21 | 241 | 7.2 | 1.58 | 3.19 | 7 | 44 | 49 |
| Z5 | 20 | 240 | 5.2 | 0.08 | 0.15 | 9 | 48 | 49 |
| Z6 | 20 | 239 | 4.4 | 0.36 | 1.19 | 9 | 68 | 73 |
| A6 | 21 | 238 | 3.9 | 0.17 | 0.06 | 8 | 54 | 59 |
| I1 | 14 | 172 | 2.2 | 0.01 | 0.03 | 25 | 12 | 13 |
| B1 | 20 | 246 | 2.0 | 0.26 | 0.19 | 7 | 35 | 45 |

To construct a dominance hierarchy, we used agonistic interactions recorded during continuous focal animal sampling and *ad libitum* sampling. An agonistic interaction was defined by the occurrence of at least one aggressive (open-mouth threat, stare, lunge, charge, chase, physical aggression) and/or submissive (make room, give ground, flee) behavior. Aggressive and submissive behaviors occurring in quick succession (<20 seconds) of each other were considered as part of the same conflict. We calculated one dominance hierarchy for the entire study period using normalized David’s score (de Vries et al., 2006) in R (version 3.6.2; R Core Team, 2019) using the function DS from the package EloRating [version 0.46.8], generating a continuous measure of individual winning success. A higher David’s score indicates higher dominance rank. The dominance hierarchy was stable throughout the whole study period.

### 2.3 Urine sample collection

Urine samples were collected from focal males whenever possible without disturbing the animal by catching it with a plastic bag or otherwise from leaves, plant litter or ground, using a disposable plastic pipette to transfer the sample to 2 ml cryotubes. When pipetting was not possible we absorbed urine using a salivette (Salivette Cortisol, Sarstedt, Nümbrecht, Germany; Büttler et al., 2018). Samples were only collected if they were not contaminated with fecal matter or urine from other individuals. Both samples in cryotubes and salivettes were stored in a thermos container filled with ice while in the field. At the end of the day, samples collected by salivette were centrifuged at 1500 rpm for five minutes to extract the urine, which was afterwards transferred to 2 ml cryotubes. All samples were then stored in a freezer at −20°C. At the end of the field season samples were transported on dry ice to the University of Göttingen where they were stored again at −20°C until analysis.

We determined the steroid excretion window in macaque urine based on previous radiometabolism studies (Bahr et al., 2000; Möhle et al., 2002). For cortisol, we considered behaviors that occurred within the same day (up to 13.5 hours) prior to sample collection to affect urinary hormone levels (Bahr et al., 2000). For testosterone, levels were at their highest in the urine up to six hours after injection, although they still remained somewhat elevated for up to one day (around 13.5 hours; Möhle et al., 2002). Therefore, we considered behaviors that occurred up to six hours prior to sample collection to affect urinary testosterone levels and used this time window in the main analysis of the study. Nevertheless, to determine the robustness of results we conducted additional analyses using a 13.5 hour clearance window and present these results in the supplementary material (Table S1a and Table S1b). The results were similar regardless of which excretion window was used. We excluded samples from which we had < 2 hours of observation time prior to sample collection for both testosterone and cortisol (N = 60), leaving a total of 650 urine samples for cortisol analysis. Out of these 650 urine samples, 78 (12%) were collected with salivettes [Salivette Cortisol, Sarstedt, Nümbrecht, Germany]. Samples collected by salivette were excluded from testosterone analyses but not cortisol analyses as they were validated for the former and not the latter (Büttler et al., 2018), leaving a total of 572 samples. To determine the robustness of results, we present cortisol analysis with salivette samples excluded in the supplementary material (Table S2a and Table S2b). Results including or excluding samples collected via sallivete were similar. Table 1 shows descriptive information and sample sizes for each study subject.

### 2.4 Hormone analysis

The measurement of immunoreactive cortisol (iuC) and testosterone (iuT) from the urine of male Barbary macaques has been previously validated (Rincon et al., 2019). Furthermore, we found a negative correlation between cortisol and time of day, thus further biologically validating our cortisol measurement (Table 3). For the accurate quantification of testosterone, but not cortisol, urine samples needed to be enzymatically hydrolyzed prior to hormone measurement (Rincon et al., 2019). Hydrolysis was carried out by mixing 20 - 100 µl urine of each sample with 40 µl β-Glucuronidase (K12 strain *Escherichia coli*, Prod. No. BGALS-RO, Sigma-Aldrich) and 900 µl phosphate buffer (composed of 6.0 g NaH_2_PO_4_ x H_2_O + 14.5 g Na_2_HPO_4_ x 2 H_2_O dissolved in 500 ml water; pH 6.9) and incubating in a shaking water bath at 37°C overnight. Hydrolyzed samples were then purified and extracted using solid phase extraction cartridges (Chromabond HR-X, 30mg, 1ml, Macherey-Nagel, Dueren, Germany) placed onto a 12-port vacuum manifold. Prior to sample loading, cartridges were conditioned first with 1 ml of MeOH, followed by 1 ml H_2_O and 1 ml phosphate buffer. Columns were not allowed to run dry at this step. Hydrolyzed urine samples were then pipetted into pre-labelled cartridges and the solution was let to sink in. Cartridges were subsequently rinsed three times with 1 ml H_2_O followed by two times with 1 ml MeOH/H_2_O (40/60) solution, after which cartridges were dried by applying a vacuum. Steroids were finally eluted with 1 ml absolute MeOH followed by 1 ml ethyl acetate, and eluates were then evaporated to dryness at 40°C under pressurized air. Then, 1 ml hydrolysis phosphate buffer was added to the dried sample eluates and vortexed for 5 min. Afterwards 150 µl 10% K_2_CO_3_ buffer and 5 ml tert. butyl methyl ether (TBME) was added to each tube for extraction of the liberated steroids. Tubes were sealed with caps and parafilm, vortexed for 10 min on a multi-tube vortexer, then centrifuged for 5 min at 2000 rpm and subsequently stored at −20°C until the aqueous layer had frozen out. The ether phase was then poured into new collection tubes, the ether was evaporated to around 1 ml, and briefly vortexed to concentrate steroids in the bottom of the tube, and finally evaporated to dryness. Dried samples were reconstituted in 1 ml absolute MeOH by vortexing for 10 min. Extracts were then transferred into 2 ml eppendorf safe-lock tubes and stored at −20°C until analysis for testosterone concentrations. The efficiency of the combined hydrolysis and extraction procedure was assessed using internal controls of testosterone glucuronide run together with each set of samples as previously described (Rincon et al., 2019). The combined hydrolysis/extraction efficiency was 62.1 ± 4.2% mean ± SD (range: 50-70%; N = 36).

**Table 2:** Immunorective urinary testosterone levels (iuT) in relation to the frequency of triadic interactions and duration of male infant care (model 1). Male identity was included as a random effect, N subjects = 14, N urine samples = 572. CI = 95% credible intervals, Pr = proportion of the posterior samples that fall on the same side of 0 as the mean. The complementary model with results for triadic interactions between-subjects is provided in the supplementary material (Table S3) and estimates are also provided in the text.

|  | Estimate | SD | CI lower | CI upper | Pr |
| --- | --- | --- | --- | --- | --- |
| Intercept | 3.06 | 0.09 | 2.88 | 3.22 | 1.00 |
| **Test predictors** |  |  |  |  |  |
| Triadic interactions (within subjects) | -0.18 | 0.07 | -0.33 | -0.06 | >0.99 |
| Infant care (within subjects) | 0.07 | 0.05 | -0.04 | 0.17 | 0.92 |
| Infant care (between subjects) | 0.08 | 0.09 | -0.10 | 0.26 | 0.82 |
| **Control predictors** |  |  |  |  |  |
| Aggression | -0.03 | 0.04 | -0.10 | 0.04 | 0.78 |
| Dominance rank | -0.16 | 0.08 | -0.32 | 0.00 | 0.97 |
| Grooming | 0.07 | 0.04 | -0.01 | 0.14 | 0.96 |
| Daytime | -0.01 | 0.04 | -0.10 | 0.08 | 0.63 |

**Table 3:** Immunoreactive urinary cortisol levels (iuC) in relation to the frequency of triadic interactions and duration of male infant care (model 2). Male identity was included as a random effect, N subjects = 14, N urine samples = 650. CI = 95% credible intervals, Pr = proportion of the posterior samples that fall on the same side of 0 as the mean. The complementary model with results for triadic interactions between-subjects is provided in the supplementary material (Table S4) and estimates are also provided in the text.

|  | **Estimate** | **SD** | **CI lower** | **CI upper** | **Pr** |
| --- | --- | --- | --- | --- | --- |
| Intercept | 5.15 | 0.14 | 4.86 | 5.39 | 1.00 |
| **Test predictors** |  |  |  |  |  |
| Triadic interactions (within subjects) | -0.24 | 0.11 | -0.47 | -0.04 | 0.99 |
| Infant care (within subjects) | 0.07 | 0.05 | -0.03 | 0.17 | 0.93 |
| Infant care (between subjects) | -0.14 | 0.13 | -0.41 | 0.12 | 0.88 |
| **Control predictors** |  |  |  |  |  |
| Aggression | -0.03 | 0.04 | -0.11 | 0.03 | 0.85 |
| Dominance rank | -0.17 | 0.11 | -0.38 | 0.04 | 0.95 |
| Grooming | 0.00 | 0.06 | -0.12 | 0.11 | 0.51 |
| Daytime | -0.23 | 0.04 | -0.31 | -0.15 | 1.00 |

We measured iuT from samples following extraction and hydrolysis (described above), and iuC from unprocessed diluted urine by using enzyme immunoassays as described in detail in Rincon et al. (2019). For the testosterone assay (Palme and Möstl, 1994), sensitivity at 90% binding was 0.3 pg; intra-assay CVs of high and low value quality controls were <10%, while figures for inter-assay CVs were 13.2% (high) and 19.5% (low). For the cortisol assay (Palme and Möstl, 1997), sensitivity at 90% binding was 0.6 pg; intra-assay CVs of high and low value quality controls were <10%, while figures for inter-assay CVs were 9.2% (high) and 10.1% (low). To account for differences in the volume and concentration across urine samples, iuT and iuC levels were corrected for creatinine and are expressed as ng/mg Cr (Bahr et al., 2000).

### 2.5 Statistical analyses

To test whether triadic interactions were associated with urinary testosterone and cortisol levels, we fitted Bayesian multilevel linear regression models with a Gaussian response distribution and identity link function. Immunoreactive urinary testosterone (iuT; model 1) and cortisol (iuC; model 2) were our response variables. Distributions of both hormone values had a strong positive skew so we used a natural log transformation to achieve more symmetrical distributions and used these values in analysis. Our test predictors were count of triadic interactions and duration of male infant care. We included duration of grooming between two adults (male-female or male-male), count of male-male aggression, a continuous measure of dominance rank (normalized David’s score), and time of sample collection (daytime) as control predictors in both models. Counts of triadic interactions and male-male aggression, and duration of infant care and grooming were calculated for the hormone excretion window of each urine sample and corrected for the observation time of the hormone excretion window. Each male had one dominance rank score for the entire study period (Table 1). All predictors were z-transformed to a mean of 0 and standard deviation of 1 to improve model convergence and interpretation of results (Schielzeth, 2010).

To decouple whether triadic interactions and infant care influenced hormone levels within- or between-subjects, we computed within-subjects centering as described in van de Pol and Wright (2009). We first calculated the mean rates of triadic interactions for each male. Then we calculated relative rates of triadic interactions by subtracting a male’s mean rate of interactions from the rate at a given observation. Therefore, relative rates of triadic interactions are centered around zero, with negative and positive values denoting deviations from the mean. The mean rates of triadic interactions were included in the models as an expression of between-subjects variation and the relative rates of triadic interactions at a given observation were included as an expression of within-subjects variation. This procedure was repeated analogously for infant care. Mean rates of triadic interactions were strongly correlated with the mean duration of infant care (r = 0.90 in the testosterone excretion window, r = 0.88 in the cortisol excretion window), making it difficult to distinguish independent effects for the between-subjects effect of each behavior. Therefore, we decided not to include both predictors within the same model to avoid issues of collinearity and instead fit two versions of the same model: one model included the mean duration of infant care and one the other included mean rates of triadic interactions. Triadic interactions and infant care within-subjects were not strongly correlated and therefore could be included in the same model. Here we report models including mean duration of infant care (and excluding mean rates of triadic interactions) and include results of the complementary model (that includes mean rates of triadic interactions but excludes mean duration of infant care) in the supplementary material (Table S3, Table S4). Regardless of which behavior was included, model estimates for all predictors remained remarkably similar and thus did not change the interpretation of results.

We fitted models in R (version 3.6.2; R Core Team, 2019) using the function brm from the package brms (version 2.11.1; Bürkner, 2017). The package brms calls on the computational framework Stan (https://mc-stan.org) to fit Bayesian models (Bürkner, 2017). We included male identity as a random effect in all models to control for multiple observations (urine samples) per subject. We also included random slopes and correlation parameters between random intercepts and random slopes for all predictors that varied within-subjects (triadic interactions within-subjects, infant care within-subjects, grooming, aggression, daytime) (Barr et al., 2013). We ran all models with 4000 iterations over four MCMC chains, which included 1000 “warm up” iterations for each chain, resulting in a total of 12000 posterior samples (Bürkner, 2017). In all models, there were no divergent transitions during warm up, all Rhat values were equal to 1.00, and visual inspection of a plot of the chains indicated that the models were able to converge. We used weakly informative priors to improve convergence, guard against overfitting, and regularize parameter estimates (McElreath, 2016). As a prior for the intercept and beta coefficients we used a normal distribution with a mean of 0 and a standard deviation of 1; for the standard deviation of group level (random) effects and sigma we used a Half-Cauchy distribution with location 0 and scale parameter 1; for the correlation between random slopes we used LKJ Cholesky prior with eta 2.

We report model estimates as the mean of the posterior distribution with 95% credible intervals (CI). To aid in the interpretation of whether predictor variables substantially affected the response (hormone levels), we calculated the proportion (Pr) of posterior samples that fell on the same side of 0 as the mean. The Pr ranges from 0.5 to 1.0, with Pr = 1.00 indicating that, given the model, the effect of a predictor was entirely positive or negative, whereas Pr = 0.5 indicates that the effect was centered around 0.

The data and code to reproduce the analyses in this paper are available at https://osf.io/4dwgu/?view_only=732edeb15854463983a00ecf5abdb18d.

## 3 Results

We recorded a total of 1701 triadic interactions during 3289 hours of observation in one non-mating season. All males of the study group were observed to partake in triadic interactions at least once (Table 1). The majority (64%) of triadic interactions involved the one newborn male, 27% involved yearlings (two males and one female) and 9% involved older juveniles up to four years of age. Rates of triadic interactions were negatively correlated with iuT levels within-subjects (Table 2; Fig. 1 *a, b*), but not between-subjects (Estimate = 0.10, 95% CI: −0.07, 0.26, Pr = 0.88; Table S3). Likewise, rates of triadic interactions were negatively correlated with iuC levels within-subjects (Table 3; Fig. 2, *a, b*), but not between-subjects (Estimate = −0.14, 95% CI: −0.40, 0.10, Pr = 0.88; Table S4).

**Fig. 1:**
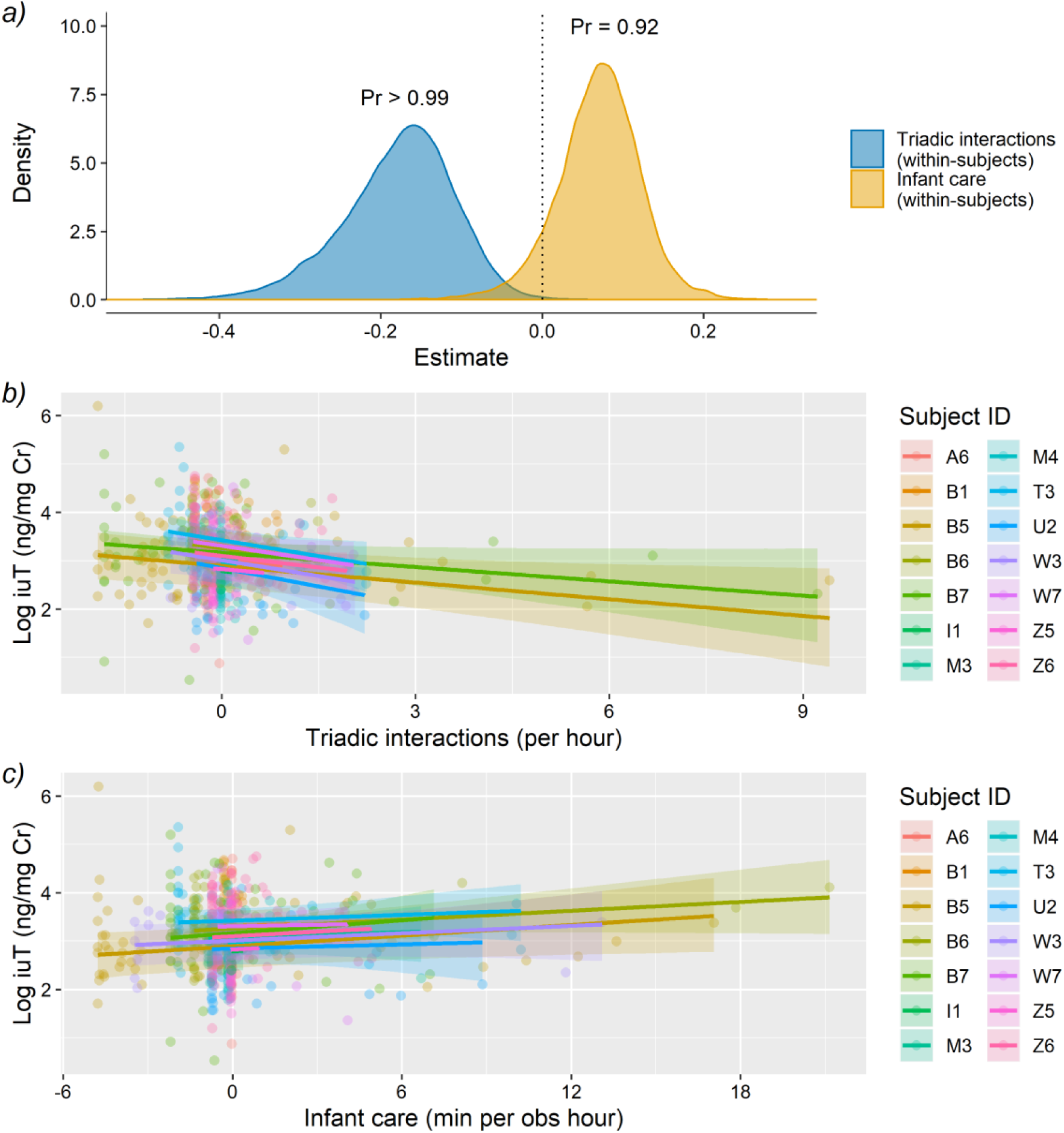
Immunoreactive urinary testosterone levels (iuT) in relation the frequency of triadic interactions and duration of male infant care within-subjects (model 1). a) Posterior probability distribution of the slope estimate. Pr = proportion of the posterior samples that fall on the same side of 0 as the mean. b) and c) Model fitted values (solid lines) and 95% credible intervals (shaded areas) per focal male, when all other predictors are at their mean. Note that the x-axis has been centered within-subjects, thus positive and negative values indicate deviations from the mean. Circles indicate raw data points (urine samples; N = 572). Note that multiple behaviors can occur within the excretion window per urine sample and each behavior can influence hormone levels in opposing directions. The individual effects of each behavior are controlled for in the full statistical model. Cr = creatinine.

**Fig. 2:**
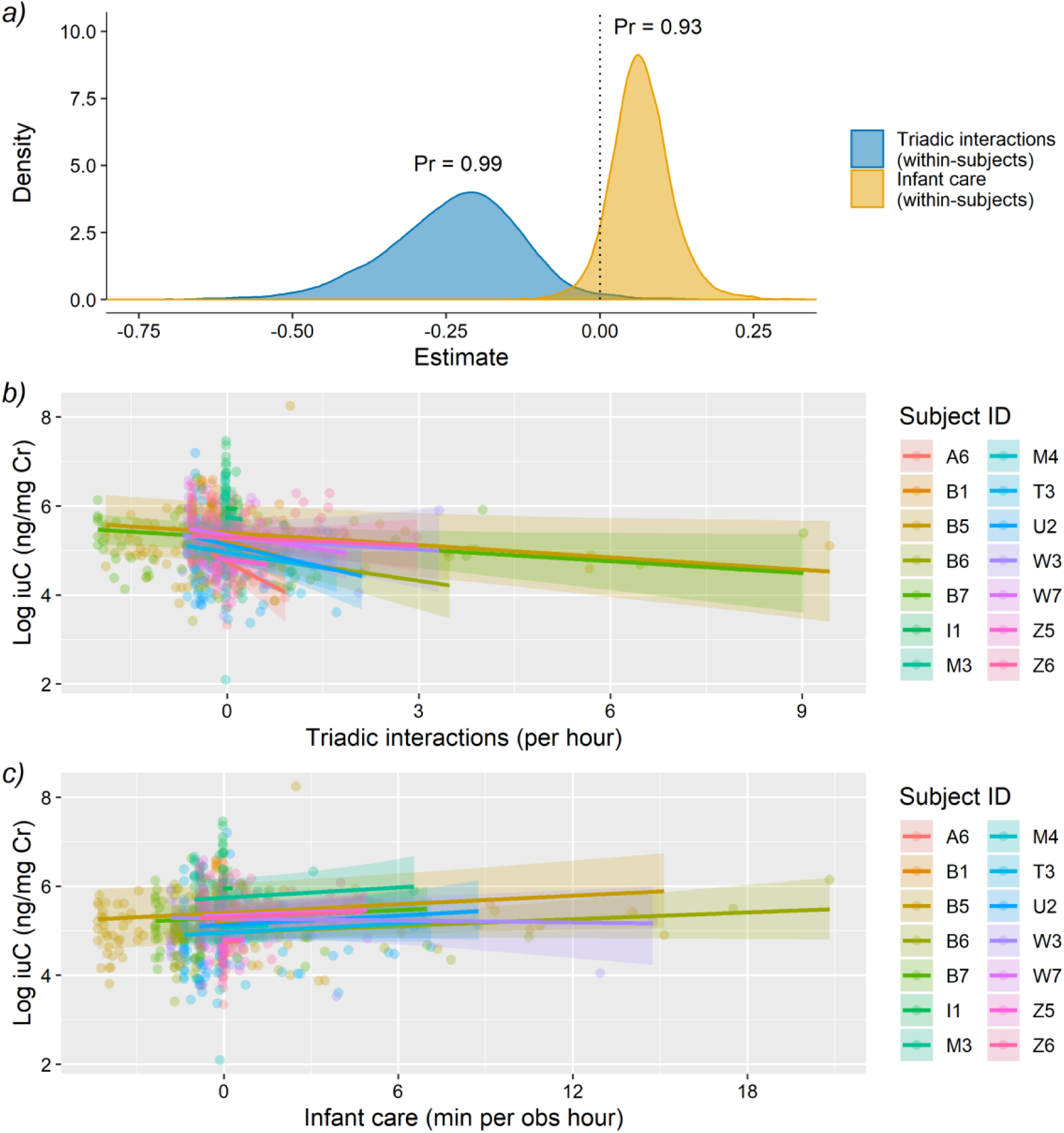
Immunoreactive urinary cortisol levels (iuC) in relation to the frequency of triadic interactions and duration of male infant care within-subjects (model 2). a) Posterior probability distribution of the slope estimate. Pr = proportion of the posterior samples that fall on the same side of 0 as the mean. b) and c) Model fitted values (solid lines) and 95% credible intervals (shaded areas) per focal male, when all other predictors are at their mean. Note that the x-axis has been centered within-subjects, thus positive and negative values indicate deviations from the mean. Circles indicate raw data points (urine samples; N = 650). Note that multiple behaviors can occur within the excretion window per urine sample and each behavior can influence hormone levels in opposing directions. The individual effects of each behavior are controlled for in the full statistical model. Cr = creatinine.

All adult males of the group were observed caring for the newborn infant and/or yearlings (≤ 1.5 years old), including time spent huddling, carrying, or grooming (Table 1). Male infant care tended to be positively correlated with iuT levels within-subjects (Table 2; Fig. 1, *a, c*), but was not related to iuT levels between-subjects (Table 2). Male infant care tended to be positively correlated with iuC levels within-subjects (Table 3; Fig. 2, *a, c*), but was not related to iuC levels between-subjects (Table 3).

## 4 Discussion

In this study, we investigated how testosterone and cortisol levels were related to social bonding and nurturing in male Barbary macaques. We first tested for a relationship between testosterone and triadic male-infant-male interactions, a ritualized behavior hypothesized to promote and/or maintain social bonds in male macaques (Berghänel et al., 2011; Kalbitz et al., 2017; Paul et al., 1996). We found that iuT was negatively correlated to frequency of triadic interactions within-subjects but was not correlated to mean rates of interactions between-subjects. In other words, the more frequently a male engaged in this behavior, the lower his iuT levels. This finding supports the S/P theory, which proposes that low testosterone levels are preferred during bonding behaviors, so as to not inhibit the bonding process (van Anders et al., 2011). These results are also in line with a previous study in humans that found decreases in testosterone levels from baseline following a friendship formation task between same-sex adults (Ketay et al., 2017). Similarly, chimpanzee males that participated in meat-sharing after a hunt - a bond promoting behavior - had lower testosterone than males that did not hunt or share, although it is unclear whether this resulted from differences in baseline levels or reactivity (Sobolewski et al., 2012). Same-sex adults may see each other as reproductive rivals and high testosterone levels may consequently interfere with the bonding process by cognitively priming individuals to be competitive rather than nurturing (Boksem et al., 2013; Eisenegger et al., 2011). Human, chimpanzee, and macaque males readily form close social bonds and long-term cooperative relationships with a few co-resident males that compete with the other co-residents (Aktipis et al., 2018; Schülke et al., 2010; Watts, 2002; Young et al., 2013). Thus, males may have been under selective pressure to downregulate testosterone during periods of affiliation to facilitate bond formation. It would be interesting to test whether lowered testosterone after affiliation between same-sex dyads is primarily found in species where long-term cooperative relationships increase fitness or if low testosterone is also found after opportunistic short-term affiliation. Beyond the formation of same-sex social bonds, high testosterone levels may also be detrimental to the maintenance of these bonds in the long term. For example, territory holder male manakins (*Pipra filicauda*, a lekking bird species), must perform cooperative displays with floater males to attract females; however, territory holders with high testosterone levels fail to maintain stable display partnerships with floater males, thus reducing their ability to compete for females (Ryder et al., 2020).

We additionally found that iuC levels were negatively correlated within-subjects with rates of triadic interactions in male Barbary macaques. Triadic interactions decrease tensions between adult males, as per the agonistic buffering hypothesis (Deag and Crook, 1971; Paul et al., 1996), and thus may consequently reduce cortisol and testosterone levels. Another non-mutually exclusive explanation relates to the potential bond-formation properties of triadic interactions, resulting in integration into the male social network (Henkel et al., 2010) and increased social support (Berghänel et al., 2011). Social support is a powerful regulator of the HPA axis and thus of glucocorticoid levels (Hostinar et al., 2014). Merely the presence of closely bonded conspecifics during the occurrence of a stressor is enough to buffer HPA axis activity in a variety of species (Hostinar et al., 2014). Indeed, male Barbary macaques with strong social bonds show attenuated glucocorticoid responses to social and environmental stressors (Young et al., 2014a). Having reliable social support may also help to downregulate HPA activity even in the immediate absence of stressors (Rosal et al., 2004; Wittig et al., 2016). Thus, study males of this study may have perceived themselves as having strengthened their social bonds and ability to call on support if needed on days when they frequently engaged in triadic interactions (Berghänel et al., 2011), consequently lowering cortisol.

While lowered testosterone levels may be necessary to prevent the inhibition of bonding, the process of bonding itself is likely driven by other hormones. For instance, the S/P theory predicts that high oxytocin levels are needed to promote social bonding (van Anders et al., 2011). In a previous study on the same group of Barbary macaques we did not find an increase in oxytocin levels following triadic interactions in general but found that oxytocin levels were only higher after triadic interactions with non-bond partners (Rincon et al., 2020). If the combination of low testosterone and high oxytocin is responsible for bonding, then triadic interactions may function to form bonds selectively between non-bonded partners while physiologically not affecting existing bonds between strongly bonded partners. In this interpretation, triadic interactions are a tool for bond formation rather than bond maintenance, although, low testosterone levels after interactions with bond partners may still be beneficial to avoid damaging existing relationships (Ryder et al., 2020), and perhaps to not inhibit friendly interactions altogether. In our study subjects, 45% of triadic interactions occurred with non-bond partners (Rincon et al., unpublished data) indicating that there is substantial variation in partner identity and thus the potential to bond with previously non-bonded partners is high. Finally, the low levels of testosterone associated with triadic interactions occur independently of infant care, suggesting that it is not merely the presence of an immature individual that lowers testosterone or cortisol levels, but that the triadic interaction itself is what is salient.

Low cortisol in conjunction with low testosterone may act synergistically to reduce tension and facilitate bonding. During a dyadic friendship formation task in humans, participants desired to be closer to their partners if their partners had low cortisol levels (Ketay et al., 2017), suggesting that affiliative interactions in a relaxed state are beneficial for both partners. While low testosterone and low cortisol may be beneficial to bond formation between same-sex adults, this pattern may not generalize to other types of dyads or contexts. For example, our findings are in contrast to a previous study in humans where testosterone levels were negatively related to friendship formation and positively related to friendship maintenance within a social network, whereas the opposite was true for cortisol levels (Kornienko et al., 2016). One difference between our study and Kornienko et al. (2016) is that the social network comprised a mixed-sex group in their study.

Steroid dynamics in relation to opposite-sex bonding may differ from that for same-sex bonding as opposite-sex dyads having greater potential for a sexual relationship and same-sex dyads having greater potential to be reproductive competitors. Indeed, high testosterone levels most likely promotes males to affiliate with females and facilitates the initial stage of sexual or romantic relationships (Goymann et al., 2019; Roney and Gettler, 2015; van Anders et al., 2011). Although high testosterone levels may be detrimental to satisfaction in long-term romantic relationships (Roney and Gettler, 2015), it is unclear if this is also the case for long-term opposite-sex platonic friendships.

We found that male infant care was weakly positively correlated with testosterone levels within-subjects but not between-subjects. Cortisol exhibited the same pattern, which is in line with a previous finding in this species (Henkel et al., 2010). These results suggest that, from the male’s perspective, infant care is neither nurturing nor relaxing and may instead be performed under a competitive context. Across many mammals, infants are at risk of infanticide in species where the period of lactation is long relative to gestation (van Schaik, 2000), as is the case in primates. Infant caretakers perform a protective role in the majority of Old World nonhuman primates and the threat of infanticide has been shown to elicit elevated testosterone and glucocorticoid responses (Cheney et al., 2015; Muller, 2017). Beyond defense against infanticide, fathers support or tolerance of immatures can have additional benefits. For instance, male baboons (*Papio cynocephalus*) often support their offspring in agonistic conflicts against other group members, which may help in gaining dominance rank and reducing stress or injury (Buchan et al., 2003). Father-offspring associations can also lead to improved feeding opportunities and ultimately faster maturation for juveniles (Huchard et al., 2013). Male Barbary macaques prefer to interact with infants based on their past mating success with the mother (Kubenova et al., 2019), which may be indicative of paternal investment. Thus, modest increases in testosterone and cortisol when males care for infants could serve to make them more alert to their social environment and more effective at providing protective support. Additionally, a positive relationship between testosterone and infant care is in line with the suggestion that infants can be used as tools for mating effort in this species (Ménard et al., 2001). It has also previously been suggested that parental effort may enhance status in young human fathers thus resulting in elevated testosterone (Mazur, 2014). It is not clear if infant care enhances status in Barbary macaque males, but it does seem likely that these behaviors are perceived positively by the infant’s mother (Ménard et al., 2001).

Differences in baseline or mean hormone levels among individuals could potentially result in different behavioral profiles where individuals perform certain behaviors at a higher or lower rate on average (Hau and Goymann, 2015). In our Barbary macaques males, however, we found that mean differences in iuT and iuC levels did not result in mean differences in rates of triadic interactions or duration of infant care. Instead, variation in steroid levels within-subjects were linked to these two behaviors. It is unclear why we found within-but not between-subjects effects given that both paternal care and friendship formation tasks influence baseline steroid levels as well as reactivity, at least in humans (e.g. Gettler et al., 2011a; Ketay et al., 2017). Differences in steroid receptor densities among male Barbary macaques could account for differences in sensitivity to hormone levels and thus result in a lack of between-subjects variation in behavior. Another possibility is that triadic interactions and infant care are expressed only sporadically and unpredictably throughout the day. Thus, selection may have acted on an individual’s ability to flexibly up- or downregulate hormone levels in response to the social environment.

## Acknowledgements

We thank Ellen Merz and Roland Hilgartner for permission to conduct the study at Affenberg Salem. We thank Lauren Cassidy, Tatjana Kaufmann and Lilah Sciaky for help with urine sample and behavioral data collection. We are grateful to Andrea Heistermann for assistance with hormone analyses. We also thank three anonymous reviewers for helpful comments that greatly improved the manuscript. This research was funded by the Deutsche Forschungsgemeinschaft (DFG, German Research Foundation) - Project number 254142454 / GRK 2070.

