## Supplementary material for "Testosterone and cortisol are negatively associated with ritualized bonding behavior in male macaques"

Table S1a: Immunorective urinary testosterone levels (iuT) in relation to the frequency of triadic interactions and duration of male infant care. This model includes results for infant care between-subjects and is complemented by Table S1b. For this model we considered a hormone clearance window of 13.5 hours, and samples collected via salivettes (N= 78) were excluded from analysis. Male identity was included as a random effect, N subjects = 14, N urine samples = 572. CI = 95% credible intervals, Pr = proportion of the posterior samples that fall on the same side of 0 as the mean.

| | **Estimate** | **SD** | **CI lower** | **CI upper** | **Pr** |
|---|---|---|---|---|---|
| Intercept | 3.06 | 0.09 | 2.87 | 3.22 | 1.00 |
| **Test predictors** | | | | | |
| Triadic interactions (within subjects) | -0.19 | 0.07 | -0.35 | -0.06 | >0.99 |
| Infant care (within subjects) | 0.08 | 0.06 | -0.05 | 0.19 | 0.93 |
| Infant care (between subjects) | 0.07 | 0.09 | -0.11 | 0.25 | 0.80 |
| **Control predictors** | | | | | |
| Aggression | -0.04 | 0.04 | -0.11 | 0.03 | 0.87 |
| Dominance rank | -0.15 | 0.08 | -0.31 | 0.00 | 0.97 |
| Grooming | 0.06 | 0.04 | -0.01 | 0.14 | 0.96 |
| Daytime | -0.00 | 0.04 | -0.09 | 0.08 | 0.51 |

Table S1b: Immunorective urinary testosterone levels (iuT) in relation to the frequency of triadic interactions and duration of male infant care. This model includes results for triadic interactons between-subjects and is complemented by Table S1a. For this model we considered a hormone clearance window of 13.5 hours, and samples collected via salivettes (N= 78) were excluded from analysis. Male identity was included as a random effect, N subjects = 14, N urine samples = 572. CI = 95% credible intervals, Pr = proportion of the posterior samples that fall on the same side of 0 as the mean.

| | **Estimate** | **SD** | **CI lower** | **CI upper** | **Pr** |
|---|---|---|---|---|---|
| Intercept | 3.06 | 0.09 | 2.87 | 3.22 | 1.00 |
| **Test predictors** | | | | | |
| Triadic interactions (between subjects) | 0.08 | 0.09 | -0.09 | 0.25 | 0.85 |
| Triadic interactions (within subjects) | -0.19 | 0.07 | -0.35 | -0.06 | >0.99 |
| Infant care (within subjects) | 0.08 | 0.06 | -0.04 | 0.19 | 0.93 |
| **Control predictors** | | | | | |
| Aggression | -0.04 | 0.04 | -0.12 | 0.03 | 0.88 |
| Dominance rank | -0.14 | 0.08 | -0.29 | 0.01 | 0.97 |
| Grooming | 0.07 | 0.04 | -0.01 | 0.14 | 0.96 |
| Daytime | -0.00 | 0.04 | -0.08 | 0.09 | 0.51 |

Table S2a: Immunorective urinary cortisol levels (iuC) in relation to the frequency of triadic interactions and duration of male infant care. This model includes results for infant care between-subjects and is complemented by Table S2b. For this model, we considered a hormone clearance window of 13.5 hours, and samples collected via salivettes (N= 78) were excluded from analysis.  Male identity was included as a random effect, N subjects = 14, N urine samples = 572. CI = 95% credible intervals, Pr = proportion of the posterior samples that fall on the same side of 0 as the mean.

| | **Estimate** | **SD** | **CI lower** | **CI upper** | **Pr** |
|---|---|---|---|---|---|
| Intercept | 5.17 | 0.13 | 4.89 | 5.40 | 1.00 |
| **Test predictors** | | | | | |
| Triadic interactions (within subjects) | -0.28 | 0.12 | -0.54 | -0.07 | 0.99 |
| Infant care (within subjects) | 0.09 | 0.05 | 0.00 | 0.19 | 0.98 |
| Infant care (between subjects) | -0.11 | 0.12 | -0.35 | 0.12 | 0.85 |
| **Control predictors** | | | | | |
| Aggression | -0.03 | 0.04 | -0.11 | 0.04 | 0.78 |
| Dominance rank | -0.17 | 0.10 | -0.38 | 0.03 | 0.96 |
| Grooming | 0.02 | 0.06 | -0.10 | 0.14 | 0.66 |
| Daytime | -0.23 | 0.04 | -0.31 | -0.14 | 1.00 |

Table S2b: Immunorective urinary cortisol levels (iuC) in relation to the frequency of triadic interactions and duration of male infant care. This model includes results for triadic interactions between-subjects and is complemented by Table S2a. For this model, we considered a hormone clearance window of 13.5 hours, and samples collected via salivettes (N= 78) were excluded from analysis.  Male identity was included as a random effect, N subjects = 14, N urine samples = 572. CI = 95% credible intervals, Pr = proportion of the posterior samples that fall on the same side of 0 as the mean.

| | **Estimate** | **SD** | **CI lower** | **CI upper** | **Pr** |
|---|---|---|---|---|---|
| Intercept | 5.16 | 0.13 | 4.86 | 5.39 | 1.00 |
| **Test predictors** | | | | | |
| Triadic interactions (between subjects) | -0.13 | 0.12 | -0.38 | 0.10 | 0.88 |
| Triadic interactions (within subjects) | -0.28 | 0.12 | -0.53 | -0.07 | 0.99 |
| Infant care (within subjects) | 0.09 | 0.05 | 0.01 | 0.19 | 0.98 |
| **Control predictors** | | | | | |
| Aggression | -0.03 | 0.04 | -0.11 | 0.04 | 0.76 |
| Dominance rank | -0.19 | 0.10 | -0.40 | 0.01 | 0.97 |
| Grooming | 0.02 | 0.06 | -0.10 | 0.14 | 0.65 |
| Daytime | -0.23 | 0.04 | -0.31 | -0.14 | 1.00 |

Table S3: Immunorective urinary testosterone levels (iuT) in relation to the frequency of triadic interactions and duration of male infant care. This is the complementary model to model 1 in the manuscript (Table 2) and includes results for triadic interactions between-subjects. Male identity was included as a random effect, N subjects = 14, N urine samples = 572. CI = 95% credible intervals, Pr = proportion of the posterior samples that fall on the same side of 0 as the mean.

| | **Estimate** | **SD** | **CI lower** | **CI upper** | **Pr** |
|---|---|---|---|---|---|
| Intercept | 3.06 | 0.08 | 2.89 | 3.22 | 1.00 |
| **Test predictors** | | | | | |
| Triadic interactions (between subjects) | 0.10 | 0.08 | -0.07 | 0.26 | 0.88 |
| Triadic interactions (within subjects) | -0.18 | 0.07 | -0.34 | -0.06 | >0.99 |
| Infant care (within subjects) | 0.07 | 0.05 | -0.04 | 0.17 | 0.92 |
| **Control predictors** | | | | | |
| Aggression | -0.03 | 0.04 | -0.10 | 0.04 | 0.79 |
| Dominance rank | -0.14 | 0.07 | -0.28 | -0.00 | 0.98 |
| Grooming | 0.07 | 0.04 | -0.01 | 0.15 | 0.97 |
| Daytime | -0.01 | 0.04 | -0.10 | 0.08 | 0.64 |

Table S4: Immunoreactive urinary cortisol levels (iuC) in relation to the frequency of triadic interactions and duration of male infant care. This is the complementary model to model 2 in the manuscript (Table 3) and includes results for triadic interactions between-subjects. Male identity was included as a random effect, N subjects = 14, N urine samples = 650. CI = 95% credible intervals, Pr = proportion of the posterior samples that fall on the same side of 0 as the mean.

| | **Estimate** | **SD** | **CI lower** | **CI upper** | **Pr** |
|---|---|---|---|---|---|
| Intercept | 5.15 | 0.13 | 4.86 | 5.38 | 1.00 |
| **Test predictors** | | | | | |
| Triadic interactions (between subjects) | -0.14 | 0.12 | -0.40 | 0.10 | 0.88 |
| Triadic interactions (within subjects) | -0.24 | 0.11 | -0.47 | -0.05 | 0.99 |
| Infant care (within subjects) | 0.07 | 0.05 | -0.02 | 0.17 | 0.93 |
| **Control predictors** | | | | | |
| Aggression | -0.03 | 0.03 | -0.11 | 0.03 | 0.85 |
| Dominance rank | -0.19 | 0.10 | -0.39 | 0.02 | 0.96 |
| Grooming | 0.00 | 0.06 | -0.11 | 0.11 | 0.51 |
| Daytime | -0.23 | 0.04 | -0.31 | -0.15 | 1.00 |
